# Sperm should evolve to make female meiosis fair

**DOI:** 10.1101/005363

**Authors:** Yaniv Brandvain, Graham Coop

**Affiliations:** Department of Plant Biology, University of Minnesota — Twin Cities. St. Paul MN, 55108; Center for Population Biology & Department of Evolution and Ecology University of California — Davis. Davis, CA, 95616

## Abstract

Genomic conflicts arise when an allele gains an evolutionary advantage at a cost to organismal fitness. Oögenesis is inherently susceptible to such conflicts because alleles compete for inclusion into the egg. Alleles that distort meiosis in their favor (i.e. meiotic drivers) often decrease organismal fitness, and therefore indirectly favor the evolution of mechanisms to suppress meiotic drive. In this light, many facets of oögenesis and gametogenesis have been interpreted as mechanisms of protection against genomic outlaws. That females of many animal species do not complete meiosis until after fertilization, appears to run counter to this interpretation, because this delay provides an opportunity for sperm-acting alleles to meddle with the outcome of female meiosis and help like alleles drive in heterozygous females. Contrary to this perceived danger, the population genetic theory presented herein suggests that, in fact, sperm nearly always evolve to increase the fairness of female meiosis in the face of genomic conflicts. These results are consistent with the apparent sperm dependence of the best characterized female meiotic drivers in animals. Rather than providing an opportunity for sperm collaboration in female meiotic drive, the ‘fertilization requirement’ indirectly protects females from meiotic drivers by providing sperm an opportunity to suppress drive.

## Introduction

Despite the apparent unity of the organism, ‘selfish’ alleles can gain an evolutionary advantage at a cost to individual fitness (Burt and Trivers, 2006), often by exploiting meiosis and gametogenesis. Because only one of the four products of female meiosis is transmitted to the egg, female meiosis is particularly vulnerable to such exploitation (Sandler and Novitski, 1957; Pardo-Manuel De Villena and Sapienza, 2001a). An allele that biases female meiosis in its favor (i.e. a meiotic driver), can increase in frequency even if it entails a pleiotropic fitness cost (Prout et al., 1973), generating a genetic conflict between the success of the driver and the organism. Meiotic drivers observed in both plants (Buckler et al., 1999; Fishman and Willis, 2005; Fishman and Saunders, 2008), and animals (Agulnik et al., 1990; Wu et al., 2005; Pardo-Manuel De Villena and Sapienza, 2001b) highlight this conflict - the selfish benefits and the associated pleiotropic fitness costs of drive sustain a balanced polymorphism (Prout et al., 1973), and often generate ongoing evolutionary escalations of drive suppressors and enhancers (Dawe and Cande, 1996; Fishman and Saunders, 2008). The threat of meiotic drive to organismal fitness is potentially so severe that many basic properties of meiosis and oogenesis, including the initial genome doubling in meiosis I (Haig and Grafen, 1991), arrested female meiosis (Mira, 1998), the structure of centromere machinery (Malik 2002 and Henikoff, 2009), and sex differences in the recombination rate (Haig, 2010; Brandvain and Coop, 2012) have perhaps evolved to enforce fairness by disrupting meiotic drive (Rice, 2013).

It is therefore somewhat surprising that despite the intense evolutionary pressure on female meiosis to prevent meiotic drive, it is potentially open to sabotage by a virtual stranger a haploid sperm genome. That is, in many animal species, female meiosis is completed only after fertilization (Masui, 1985), creating ample opportunity for interaction between the sperm and female meiotic machinery (note that, across animals the variation in timing of sperm entry into the egg and the timing at which female meiosis stalls (Figure S1) complicates this opportunity in some taxa, and that the alternation of generations likely precludes this interaction in plants). Therefore, in many species a ‘green-bearded’ (Gardner and West, 2010) sperm-acting allele that recognizes and facilitates the meiotic drive of a genetically equivalent allele in heterozygous females could presumably rapidly spread through a population. At first sight, female meiosis appears primed for conflict caused by such selfish systems. Here we ask if sperm do indeed evolve to collaborate with female drivers to exploit this apparent weakness in the defense against meiotic drive.

Before doing so, we highlight the evidence that sperm can (or do), influence female meiosis. It is becoming increasingly clear that sperm bring a wide variety of RNA and proteins into the egg (Miller et al., 2005). Some of these have known functions, for example, in most animal species, sperm – not eggs – are responsible for the transmission of the centriole, a vital component of the mitotic machinery for the zygote (Schatten, 1994). Detailed functional studies and analyses of paternal effect mutations in model systems further highlight that sperm-transmitted products have a wide-range of functions in egg activation, completion of syngamy, zygotic development, and the resumption and successful completion of female meiosis (e.g. Yasuda et al., 1995; Loppin et al., 2005; Miller et al., 2001; McNally and McNally, 2005; Churchill et al., 2003). For example, in *C. elegans*, premature deployment of the sperm aster disrupts MII meiotic segregation in the egg, leading to a triploid zygote (McNally et al., 2012). However, the function of many of the products the sperm brings into the egg is completely unknown and these products vary widely over species (Karr et al., 2009). It seems quite plausible that sperm-based products, and hence sperm haplotype or paternal genotype could influence various aspects of female meiosis that occur after fertilization.

Current evidence from the best characterized systems of female meiotic drive in animals (the *In* and *Om* loci in mice) suggests that sperm influence on female meiotic drive is not only possible, but likely. While ruling out the alternative hypothesis of early selection on zygotes in these cases is challenging (see pages 52-54 in Burt and Trivers, 2006, for comment), it appears that the extent to which *In* and *Om* distort the second female meiotic division partially depends on the genotype of the fertilizing sperm (Agulnik et al., 1993; Wu et al., 2005). The fact that the two best characterized, polymorphic systems of putative female meiotic drive systems in animals show this effect suggests that if female meiotic drive is common the role of sperm in modifying female drive will be important as well.

Numerous lines of evidence suggest that female meiotic drive is a common and important evolutionary force, and therefore the opportunity for sperm to influence female drive is likely relevant to many animals. While research to date has identified a few extreme cases of female meiotic drive in the small number model systems systematically studied (Agulnik et al., 1990; Fishman and Saunders, 2008; Hiatt and Dawe, 2003; Novitski, 1951; Pardo-Manuel De Villena and Sapienza, 2001b), rapid evolution of the basic components of the meiotic machinery (e.g. centromeres, telomeres, etc …) suggest consistent selection on female meiotic drivers and suppressors of meiotic drive in many animal species (e.g. Anderson et al., 2008, 2009; Axelsson et al., 2010; Malik, 2009). We expect that over the next decade the spread of sequencing to a range of systems will reveal many more female meiotic drive systems; however, carefully characterizing them will still remain a challenging task.

Because female meiotic drive is likely a common force with predictably negative effects on organismal fitness, and because sperm have ample opportunity to influence female drive, we develop population genetic models to address the the expected influence of sperm on female drive. We first focus on models in which ‘self-promoting’ alleles in sperm facilitate drive of like alleles during gametogenesis in heterozygous females. These models show that such sperm-acting alleles have more difficulty invading a population than do traditional meiotic drivers, and under most circumstances, cannot be maintained as a balanced polymorphism. Because self-promoting drivers are unlikely to create a sustained genomic conflict, female meiosis will have little incentive to evolve resistance to them. We then examine models in which a novel sperm-acting allele modifies the efficacy of a polymorphic meiotic driver. Such models universally reveal that sperm-acting drive modifiers are favored only if they suppress drive. These results highlight the fact that the interests of sperm and maternal genomes’ are often aligned, as both are invested in the fate of the resultant zygote (as was speculated for the *In* locus, Pomiankowski and Hurst, 1993). Thus, there is little selective benefit to females in preventing sperm to influence female meioses, and in fact, females eschewing this delay would potentially lose sperm assistance in the suppression of meiotic drivers. Given the wide-spread requirement of fertilization for the completion of female meiosis, various features of the interaction between sperm and egg may result in an equitable transfer of genetic material - wether this result is the ultimate evolutionary function of the fertilization requirement or a coincidental pleiotropic outcome is beyond the scope of this manuscript, but our intuition argues against the prior (see Discussion).

## Methods

We present deterministic one and two locus population genetic models of sperm influence on female meiotic drive to evaluate wether sperm are likely to collaborate with female meiotic drivers or to stop them.

We present six related models – three single-locus ‘pleiotropy’ models and three two-locus ‘drive-modifier’ models. **Model 1** describes a single-locus female meiotic driver. **Model 2** describes a single-locus sperm-dependent female driver – that is, an allele whose transmission in female meiosis depends on sperm haplotype. **Model 3** describes a single-locus paternal-dependent female driver – an allele whose transmission in female meiosis depends on paternal genotype. Assuming that a traditional driver segregates at its equilibrium frequency (identified in Model 1), we investigate the evolution of tightly linked (**Models 4 and 5**), and unlinked (**Model 6**) sperm-dependent modifiers of drive. In **Models 4 and 5**, we treat this two-locus system as if it consists of a single locus with three alleles: *A*, *B* and *C*, corresponding to the case when the sperm-modifier is very tightly linked to the driving (**Model 4**) or non-driving allele (**Model 5**) at the drive locus such that recombination is unexpected. To evaluate the feasibility of sperm modification of female meiotic drive, as compared to female suppression of drive, we conclude with a model of female drive suppression by an unlinked female-acting suppressor (**Model** 6′). In all cases, we assume that fitness is independent of the drive modifier.

All models include a biallelic locus (*A*/*B*) with non-driving and driving alleles in frequencies *f_A_* and *f_B_* = 1 − *f_A_*, respectively, while Models 4-6 include a drive-modifying locus. Transmission rules describing the outcomes of all matings in each model are presented in a File S1. The fitness of genotype, *g*, is sex-independent and equals 1, 1 − *h_s_*, and 1 − *s* for genotypes *AA*, *AB*, and *BB*, respectively. Genotypic frequencies equal *f_g_* for adults in the current generation, 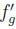 in the next generation of zygotes (i.e. after recombination, random mating, drive, and syngamy, but before selection) and 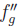 in adults after selection. After a complete generation genotype frequencies are 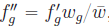, where 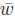 is the population mean fitness and equals 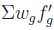.

We verbally describe our main results below. Readers interested in the details of these results should turn to the Appendix for a mathematical treatment, and to our Mathematica worksheet (File S2) for our complete derivations. There, we present critical analytical results in Equations 1−11, and describe our analyses and results in more detail. Because a number of our analyses are approximations based on assuming that genotype frequencies follow Hardy Weinberg Equilibrium (HWE), we note which analytical results are approximate. We verify these approximations with exact numerical iterations in Figure S2—S4 and File S3.

## Results

### Invasion of the population by a driving allele that promotes itself

In the standard single-locus, biallelic model of female meiotic drive, the driving allele is transmitted to the egg in heterozygotes with probability *d* > 1/2, regardless of sperm genotype (e.g. Ubeda and Haig, 2004, and see Model 1 in the Appendix for more details). To depict a case of a self-promoting meiotic driver, we modify this standard model such that the driver is only effective when fertilized by a sperm carrying that allele (see Figure 1A and Model 2 in the Appendix and File S1). We then identify the conditions allowing for the spread of this self-promoting driver, and evaluate whether a driver of this form could generate a sustained conflict favoring the evolution of suppressors. We conclude our single locus results with an analysis of a related model (Model 3) – in which drivers influence their transmission in females via paternal genotype, rather than sperm haplotype.

**Figure 1:**
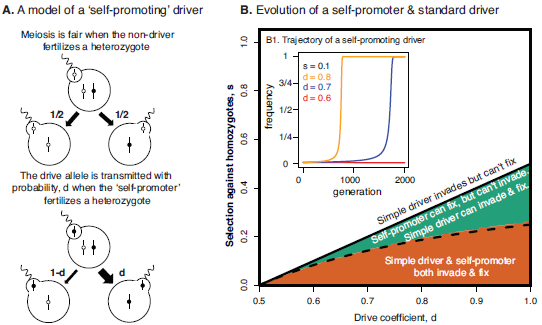
**A**. A visual depiction of our model of ‘self-promoting’ driver. Transmission probabilities for alleles through female meiosis depend on sperm genotype. The non-driving A-allele, and self-promoting B-allele are represented by unfilled and filled circles, respectively. **B**. Evolution of a self-promoter and standard driver. Assuming that the fitnesses of drive homozygotes and heterozygotes are 1 − *s* and 1, respectively. *Main figure*:Boundary conditions for the invasion and fixation of self-promoting and standard meiotic drivers, with drive coefficient, d. Colored regions depict exact results, while lines represent analytical approximations. *B*1: Trajectories of sperm-dependent female drive each allele has *s* =0.1 against the homozygotes. The drive coefficient is denoted by color.

For comparative purposes, we first briefly present the standard drive model (see e.g. Prout et al., 1973; Ubeda and Haig, 2004, for additional results). Assuming that the driving allele is deleterious in both sexes, but fully recessive (i.e. the fitness of drive homozygotes equals *W_BB_* =1 − *s* and other genotypic fitnesses equal *w_AA_* =*W_AB_* =1), it always invades because, when rare it occurs predominantly in heterozygotes and therefore drives without a fitness cost. However, when s is large (*s* > (2*d* − 1)/(2), solid black line in (Figure 1B) a driver cannot fix and will be maintained as a protected polymorphism (Prout et al., 1973). The parameter space where the allele can invade but not fix is shown in white in Figure 1B. When the allele is maintained as a polymorphism, it provides an opportunity for the evolution of drive suppressors, corresponding well to empirical examples of female meiotic drive (reviewed in Burt and Trivers, 2006).

In contrast to a traditional driver, which drives but pays effectively no fitness cost when rare, a self-promoting driver specifically creates low fitness drive homozygotes by uniting driving female gametes with sperm enabling that drive. It must therefore overcome a drive-associated homozygous fitness cost simply to spread when rare. The conditions allowing the invasion of a self-promoting driver are consequently far more restrictive than those for a standard meiotic driver. When rare, a fully recessive, self-promoting driver can only invade when s is less than approximately (2*d* − 1)/(4*d*) – see dashed black line in Figure 1B. This analytical approximation, derived from Equation (1) assuming Hardy-Weinberg, closely matches results obtained by exact numerical iteration (Figure 1B. We remind readers that Equation 1 and all equations discussed in the main text are presented in the Appendix and derived in File S2).

When a self-promoting driver does spread it spends much of its time at low frequency, because the paucity of complementary sperm compromises its ability to drive. However, once relatively common, it rapidly achieves fixation due to its positive frequency dependent behavior (Figure 1B.1). This positive frequency dependence can induce bistability in its spread - some values of s allow the fixation of this driver when common, but preclude its invasion when rare Equation 2 and Figure 1B). In this case, the driver will be fixed if its frequency exceeds some threshold (approximated in Equation 3 and presented exactly in (Figure S2) and lost otherwise. For most parameters, this threshold is likely too high to be reached by drift, and therefore the fate of a self-promoting driver is determined by the more restrictive invasion criteria rather than the fixation criteria.

Inclusion of a heterozygous fitness cost (i.e. *w_AB_* =1 − *s_h_*) further constrains the evolution of a self-promoting driver. In fact, with any heterozygous fitness cost, a rare self-promoting driver is always selected against. However, this case also displays bistability – when *s* is sufficiently small (Equation 4) this allele fixes deterministically if its frequency exceeds some threshold (Equation 5, exact results in Figure S3). This bistability prevents self-promoting drivers from invading reasonably sized populations, and assures that if they do invade, they will rapidly fix. Our model therefore predicts that self-promoting drivers will not be observed as stable polymorphisms in natural populations. This lack of a balanced polymorphism precludes the evolution of an allele that suppresses this form of meiotic drive in females. Relaxing our assumptions of panmixia by allowing for arbitrary levels of inbreeding (in the form of self-fertilization, implemented in File S3), more thoroughly aligns the interests of both parents and parental chromosomes, restricting further the possibility for invasion of both traditional female drivers and ‘self-promoting’ drivers (Figure S5). Additionally, because inbreeding reduces the frequency of heterozygotes, the invasion and fixation criteria converge, as both become stricter with increased inbreeding rates.

Although the allelic identity of sperm could plausibly influence the outcome of female meiosis, limited gene expression in sperm (e.g. Joseph and Kirkpatrick, 2004) suggests a model where sperm influence female meiosis via expression of the fertilizing male’s diploid genotype (perhaps due to proteins and RNAs packaged into the sperm), rather than sperm haplotype. This paternal-genotype dependent model (Model 3 in the Appendix) requires one additional parameter, as we exchange d in the sperm dependent case for *d_het_* and *d_hom_* which describe the transmission of the drive allele in a heterozygous female mating with males heterozygous and homozygous for the self-promoting drive allele, respectively. Here, a rare driver invades when *s* is less than (*d_het_* − 1/2)/*d_het_*, and usually fixes when it invades. However, when the distorting effect of genotype displays strong dominance in its effect on female meiosis (*d_het_* is close to *d_hom_*), a narrow sliver of parameter space sustains a polymorphism when the cost of the drive is recessive (see Figure S4, and Equation 6). While mathematically interesting, it does not seem particularly realistic to think that the effect of the drive allele would be dominant in its action through the male genotype, while the cost would be recessive. Therefore, although Model 3 can sustain a polymorphism, the lack of biological reality underlying the small portion of parameter values required for this polymorphism make us doubt its general applicability.

Given the difficulty that self-promoting meiotic drivers have entering the population, the speed at which they fix if they do, and the narrow parameter range permitting balanced polymorphisms at such loci, it seems very unlikely that such alleles could drive the evolution of female suppressors of sperm-enabled female meiotic drive.

### Two locus models of sperm-dependent female drive

Models 2 and 3, above, explored the dynamics of an allele that drove in females when signaled by a complementary signal in sperm. We complement this single-locus approach with alternative models of two loci - one a female driver, and the other, a sperm-acting allele which modifies the effect of drive upon fertilization. In this model, a female meiotic driver with no sperm-dependency initially reaches drive-viability equilibrium (with two alleles A and B are the ancestral non-driver and driver alleles, Figure 2A1). Subsequently, a sperm-acting modifier of female meiotic drive arises at another locus. In these two-locus models, the driver is transmitted to *d*_0_ of gametes from female heterozygotes when fertilized by wild-type sperm, and *d*_1_ =*d*_0_ + ϵ when fertilized by a sperm-acting drive modifier.

**Figure 2:**
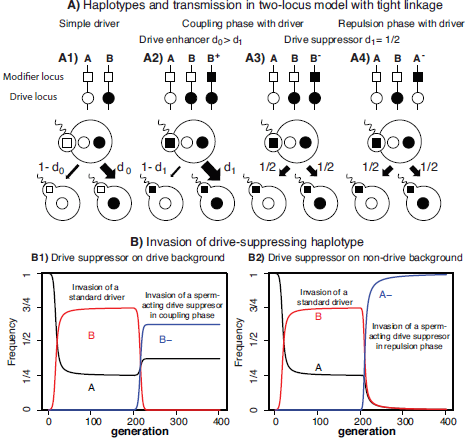
Models of a sperm-acting drive modifier tightly linked to a meiotic driver. **(A)** Sperm carrying the derived allele at the modifier locus (filled squares) alters transmission at the driving allele (filled circles) during female meiosis. Alleles at these two tightly linked loci form three haplotypes (top of A). **A1** In the standard model of drive there is no variation at the modifier, and the driver is transmitted to the egg with probability *d*_0_. **A2** The modifier allele increases the transmission of the drive allele (*d*_1_ > *d*_0_), and due to their shared genetic background, also increases its drive. **A3 & A4**) The sperm-acting modifier acts to decrease drive (*d*_1_ =1/2 in A3 & A4, or more generally, *d*_1_ < *d*_0_) and arises on the same or opposite background from the driver (A3 & A4 respectively). **(B)** Invasion of a sperm-acting drive suppressor linked to a driver. After the driver (B haplotype) reaches drive selection equilibrium, we introduce a sperm acting drive modifier. We assume full drive (*D_0_* =1), a recessive lethal fitness cost to drive (*h_s_* =0, *s* =1) and that the sperm-acting modifier results in a fair meiosis. **B1)** The B allele replaces the ancestral drive haplotype, but segregates at a lower equilibrium frequency. **B2** The A^−^ allele replaces the ancestral non-driving haplotype, and in this case, removes the driver from the population.

We first assume that the modifier is tightly linked to the drive locus (effectively creating a third allele/haplotype at this locus) and arises on the drive-background. Tight linkage offers the best chance for a collaboration to evolve between a driver and a sperm-acting drive enhancer, as recombination breaks up drive haplotypes (Thomson and Feldman, 1974; Charlesworth and Hartl, 1978; Haig and Grafen, 1991). Additionally, tight linkage between female driver and sperm modifier is consistent with the nature of well characterized drive systems which are often maintained as polymorphic inversions with numerous linked modifiers Burt and Trivers (2006). We conclude by analyzing models with alternative linkage relationship between driver and drive modifier – in Model 5 the modifier arises tightly linked to the non-driving allele, and in Model 6 it is unlinked to the driver.

When the modifier of drive arises on the drive background (i.e. in coupling phase), is tightly linked to the driver, and enhances drive we label this non-recombining drive/modifier haplotype as the *B*^+^ allele. The *B*^+^ allele acts in sperm to increase the effectiveness of drive for both the *B* and *B*^+^ alleles in *AB* and *AB*^+^ heterozygotes (see Figure 2A2, and Model 4 in the Appendix and File S1). Naively, *B*^+^ may spread by capitalizing on the additional drive it causes; however, this is not the case for a few simple reasons. First, the novel *B*^+^ haplotype arises when the ancestral driver is at drive-selection balance, and therefore immediately suffers a genotypic fitness cost equivalent to the *BB* homozygote. Worse yet, a novel *B*^+^ haplotype most often helps the *B* allele drive (*B*^+^ sperm meeting *AB* eggs), because *B* is initially more common than *B*^+^. Therefore, sperm-acting drive facilitator alleles experience a profound disadvantage in this scenario, even more so than under the previous two allele model. We have found no parameter range of this three allele system that allows the sperm-acting drive facilitator *B*^+^ to invade the population (Appendix Model 4, eqn. (8). and File S2).

While sperm enhancement of a female drive cannot displace a polymorphic female driver, sperm based drive suppressors can. Imagine a sperm-acting allele that restores fairness to female meiosis arises on the drive background, creating a third allele *B*^−^(Figure 2A3, Model 4 in Appendix). This new allele still experiences female drive when fertilized by A or B sperm, but it does not drive when fertilized by another *B*^−^ so it avoids the excess formation of low fitness genotypes. This allows the *B*^−^ to displace the ancestral driver (Figure 2B1, Equation 8), and often returns to a lower equilibrium frequency than the *B* allele (likely because it surprises its own drive), further decreasing the extent of drive in the population. If this sperm-acting drive suppressor arises on the non-driving A background (i.e. in repulsion phase, creating a third allele *A*^−^, Figure 2A4, Model 5), or is unlinked to the drive locus (Model 6), it readily invades a population segregating for the drive system (Equations 9 and 10). We note that the evolution of sperm-acting drive suppressors unlinked to a driver (Model 6) is both qualitatively and quantitatively similar to the evolution of a female-acting drive suppressor (Model 6′ – compare Equations 10 and 11).

The sperm-acting drive suppressing allele lowers the frequency of the original driver (perhaps to zero), and spreads to fixation if it does not carry strong fitness costs (Figure 2B2). This result is consistent with previous work showing that drive suppressors unlinked to, or in repulsion phase with drivers usually invade polymorphic drive systems (e.g. Brandvain and Coop, 2012). Therefore, all two-locus models of sperm influence on female drive suggest that sperm will evolve to oppose female meiotic drive, and can do so as effectively (or more effectively) than female-acting drive modifiers.

## Discussion

Sexual reproduction is a high-stakes event that determines what gets transmitted to the next generation. As a consequence of this intense competition, alleles that gain a transmission advantage during reproduction can succeed evolutionarily even if they lower organismal fitness. This generates numerous conflicts including sexual conflicts between mates (Arnqvist and Rowe, 2006), and conflicts between alleles that are over-transmitted in meiosis and the organisms they injure while doing so (Burt and Trivers, 2006). Such conflicts and their resolution likely play a major role in the structure and evolution of many basic biological processes (Rice, 2013).

### Major result: Sperm evolve to enforce fairness in female meiosis

It seems that allowing sperm to influence the outcome of female meiosis would generate a confluence of these potential conflicts – sperm could actually assist an allele that distorts female meiosis. However, this is not the case. We find that an allele which acts through sperm to distort female meiosis in its favor can rarely spread through a population if it bears any cost. Additionally, when this self-promoting driver can spread, it can only rarely be maintained as a protected polymorphism, and due to its positive frequency dependence, it spends very little time at intermediate frequency. As such, this type of exploitation cannot generate a sustained genetic conflict. It is therefore unlikely that female oogenesis and meiosis will evolve to prevent their effect. Thus, females can delay the completion of meiosis until after fertilization without risking exploitation by collaborations between female drivers and sperm alleles. Although the fertilization requirement allows sperm an opportunity to enforce fairness in female meiosis, this is unlikely it evolutionary raison d’être. In fact, to suggest so, presupposes that sperm have an evolved system, to prevent meiotic drive before they have a mechanism to do so.

### Explaining why sperm evolve to enforce fairness in female meiosis

Why is it that an allele that biases female meiosis in its favor can generate a genetic conflict, but an allele in sperm that assists this female driver cannot? So long as the transmission advantage of female meiotic drive outweighs the organismal fitness cost to heterozygotes, the female driver can spread when rare, and it increases in frequency until the fitness cost to homozygotes balances the transmission advantage. By contrast, a sperm promoter of female drive is only effective when matched with a heterozygote female – meaning that, when rare, this allele rarely enhances female drive. Even worse, when it does so it will preferentially find itself in a low fitness drive homozygote. Not only are drive-promoting sperm alleles unable to create a sustained genetic conflict, but alleles in sperm with the opposite effect – that is those that prevent their own drive through female meiosis do maintain a polymorphism and provide evolution with time and opportunity to further minimize drive. This is because such drive suppressing alleles reduced their chances of forming low fitness homozygotes. More generally, natural selection favors alleles that act through sperm to reduce the opportunity of female meiotic drive regardless of linkage or phase.

### Predictions from theory

The theory developed above has one overarching conclusion – that when possible, males evolve to make female meiosis fair. This simple result provides numerous novel predictions, many of which are directly testable.

Our most direct prediction is that for organisms in which female meiosis is not completed until after fertilization, sperm will act to suppress female drive at the stage at which they can influence meiosis. This prediction, which holds when modifier and driver are the same gene (Model 2) or are in tight linkage (Model 4), is strongly supported by the observation that female meiosis is fairer when fertilized by sperm bearing the drive allele in two of the best described cases of female meiotic drive in animals (the *Om* and *In* loci in mice, Agulnik et al., 1993; Wu et al., 2005). Both this prediction, and the empirical support for it run contrary to expectations of a naive verbal “green-beard” model.

Our model of a sperm-acting drive suppressor unlinked to a female driver (Model 6) also predicts that sperm should evolve to prevent meiotic drive; however, it contains no simple mechanism to maintain polymorphism for sperm-acting drive suppression. Given the benefit to sperm of hampering female drive, drive-suppressing sperm are often likely to be fixed within a species, making the hypothesis of sperm-acting drive-suppression difficult to test from intra-population crosses. However, crosses between populations or species are likely to provide critical tests of our theory - specifically we predict that female meiosis will be less fair when a species (or population) is fertilized by heterospecific sperm because either such sperm have not evolved to counter novel female meiotic drivers, or because antagonistic coevolution between a driver-suppressor pair has been independent since two populations have separated. We can therefore predict that segregation in F1 females backcrossed to parental species will likely be biased, with a deficit of transmission of the paternal species allele from the F1 female. These predictions follow straightforwardly from the theory presented above; however, we caution that tests of meiotic drive, and especially sperm-dependent meiotic drive require a high standard of evidence to exclude plausible alternative hypotheses such as genotypic inviability including epistatic maternal by zygotic lethality (e.g. Sawamura et al., 1993).

Our theory also encourages phylogenetic hypotheses concerning the relationship between the opportunity for female meiotic drive and the requirement of fertilization for the completion of female meiosis.

For example, we predict that a lower opportunity for female meiotic drive, e.g. an animal lineage with a history of high inbreeding or selfing, may be accompanied by a relaxation of the requirement of fertilization for the completion of female meiosis (although opportunities to test this hypothesis may be limited because lineages may only persist for a short time). This prediction follows from the logic that although the benefit of sperm protection from drivers did not necessarily favor the evolution of the fertilization requirement, mutants who forge this requirement will experience a higher level of meiotic drive than individuals who do not. Therefore removing this requirement is safest in populations with little drive. We caution that other constraints on the fertilization requirement could prevent species from conforming to this prediction.

Our results also suggests that phylogenetic variation in the stage of female meiosis when fertilization occurs (see Figure S1) may influence the prevalence of female meiotic drive. For example, centromeric drive may be more common in taxa where females complete MI before fertilization, as compared to species in which sperm interact with eggs arrested in MI, because in the prior case, sperm-based modifiers can only intercede during the second, but not the first meiotic division. As a potential test of this hypothesis, the speed of centromere turnover could be compared in species in which sperm interact with eggs paused at MI and MII (assuming the pace of centromere turnover serves as a proxy for the frequency of MI drivers).

## Conclusion

Our results highlight potentially counterintuitive results of complex genetic conflicts. Despite much opportunity for conflict between sperm and females over fertilization (Partridge and Hurst, 1998), the interests of fertilizing sperm and female are quite well aligned during syngamy. While conflict between mother and her alternative chromosomes ensues, fertilizing sperm decidedly side with mom, as both have a shared interest in producing a viable and potentially fit offspring. Our model does not directly speak to the evolutionary origin of female meiotic arrest (for a review and evaluation of such hypotheses see Mira, 1998), in fact, we presuppose its existence. However, given the existence of female meiotic arrest, and that its timing and mechanistic details are variable across species (Figure S1, and Masui. 1985; Karr et al., 2009) the nature of the meiotic arrest and interactions between sperm and egg may be molded by selection to reduce the opportunity for female meiotic drive, and counteracted by selfish drivers evolving to overcome these adaptations.

## Acknowledgements

We would like to thank members of the Coop lab for comments. This work was made possible by the NSF Postdoctoral Fellowship in Biology FY 2010 project number 1002942 awarded to Y.B..

## Models 1-3. Single-locus drive

### Model 1. Traditional driver

In the standard female drive model, meiosis in males is fair such that *A*/*B* heterozygotes contribute A and B alleles with equal probabilities; however, *A*/*B* females transmit the *B* allele with probability *d* > 1/2. We note that the timing of fertilization relative to female meiosis places another constraint on *d*, for example, if fertilization (and therefore, sperm dependent drive) takes place at MII (as in mammals), female drive requires an uneven number of crossovers between the centromere and the drive locus, so *d* is bounded to be < 0.83 (see Buckler et al., 1999, for discussion). After drive and random mating, genotype frequencies are

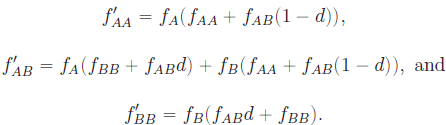

As detailed above, exact frequencies after drive, random mating and selection are 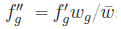 Assuming HWE, a rare driver will spread when 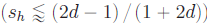, and will fix when 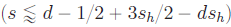. This later inequality reduces to 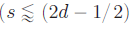 when the cost of drive is fully recessive.

### Model 2. Single locus, sperm-dependent drive

Our single-locus model of sperm-dependent drive resembles the traditional driver, with the caveat that the *B* allele drives in heterozygous females only when fertilized by *B*-bearing sperm. Therefore, genotype frequencies after drive are

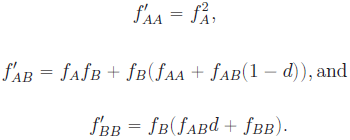

We iterate exact genotype frequency recursions 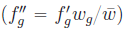 over generations to produce the frequency trajectories shown in the inset of Figure 1B by plotting 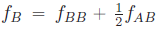 over time. To assess invasion or fixation criteria, as well as bistability points, we iterate this system and test whether *f_B_* increases over a grid of parameters.

#### Recessive fitness cost of self-promoting driver

When fully recessive, the change in frequency of the self-promoting driver across generations equals

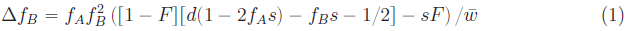

where *F* is the deviation from genotypic frequencies expected under Hardy-Weinberg. Assuming HWE (*F* =0) a common, recessive, self-promoting driver invades if 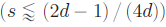, and fixes if 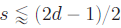. Therefore, when

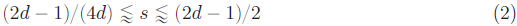

a recessive, self-promoting driver will deterministically fix if drift, mutation, or migration pressure bring its frequency above but it will be lost when introduced below this frequency. Compared to exact results (Figure S2), Equations (2) and (3) offer reasonable, but imperfect approximations.

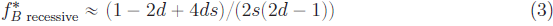

#### Cost of driver in heterozygotes

When the fitness of drive heterozygotes is compromised (*s_h_* > 0), a self-promoting driver cannot invade when rare. This results from the fact that, when rare, *B*-bearing sperm and heterozygous eggs will rarely encounter one another 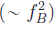) but the allele still pays a cost in heterozygous individuals (∼ *f_B_*). However, this system too, is bistable - as the driver increases in frequency it is more often fertilized by a driving sperm and therefore drives more effectively. Therefore, assuming HWE, if

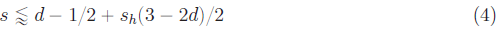

this self-promoting driver deterministically fixes when its frequency is greater than

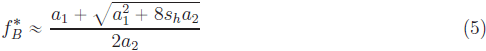

where, *a*_1_ =(1 — *2ds_h_* + 4*ds* — 2*d* — 3*s_h_*) and *a*_2_ =(2 (*s* — *s_h_*) (2*d* — 1)). Comparison of Equation 5 to exact results obtained by a simple parameter search (Figure S3) show that this approximation is reasonably correct for small parameter values; however, it underestimates 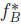 for large parameter values, presumably because they result in strong departures from HWE.

### Model 3. Single locus, paternal genotype dependent drive

In the case when female meiotic drive depends on paternal genotype, a heterozygous female will transmit the *B* allele with probabilities 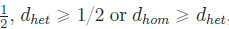, when mated with to *AA*, *AB*, or *BB* males, respectively. In this model, genotype frequencies after drive, random mating, and selection are

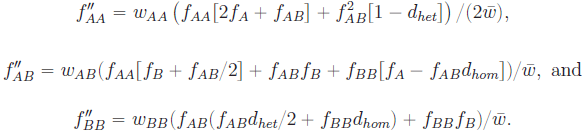

If the cost of drive is fully recessive (i.e. *s_h_* =0), assuming HWE, a rare paternal-genotype-dependent driver invades when 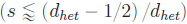, and when common, this driver fixes if 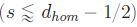, approximations well supported by exact results (Figure S4). Specifically, when drive in heterozygotes is large relative to that in homozygotes,

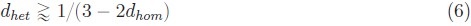

fixation criteria are more stringent than invasion criteria, and therefore some values of s can maintain a stable polymorphism. Under these parameter values, a rare paternal-genotype-dependent driver can increase in frequency because it gains a transmission advantage and suffers no fitness cost when heterozygous eggs are fertilized by A-bearing sperm of heterozygous males. As the frequency of the B allele increases, it will be unable to avoid producing unfit homozygous offspring, leaving it trapped in the population at frequency 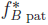. Assuming HWE, a recessive fitness cost (*h_s_* =0), and dominance of driver (*d_het_* =*d_hom_*) this equilibrium frequency is

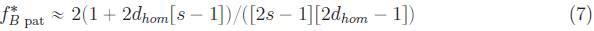

By contrast, when 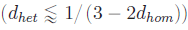 the case is reversed, and the model is bistable.

## Models 4-6. Two-locus, sperm-dependents drive

### Model 4. Drive-modifier in coupling phase

When the *C* allele is tightly linked to the driver allele, genotypic fitnesses equal *w_AC_* =*w_AB_* =1 − *s_h_*, and *w_BC_* =*w_CC_* =*w_BB_* =1 − *s*. Assuming HWE, a recessive fitness cost to drive, and assuming that the *A*/*B* locus is at its equilibrium frequency, the change in frequency of a rare drive modifier is

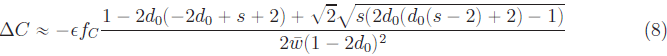

For all parameters sustaining a polymorphism at the drive locus (*s* > *d_0_* − 1/2), this corresponds to a decrease in frequency of the *C* allele when it enhances drive (ϵ > 0 - the *B*^+^ model, above), and an increase in frequency of the *C* allele when it suppresses drive (ϵ < 0 - the B^−^ model, above). More generally, even when the cost of drive is not fully recessive, the B^−^ allele will invade and fix under all parameters sustaining a previous polymorphism at the drive locus (see File S2).

### Model 5. Drive-modifier in repulsion phase

When the *C* allele is tightly linked to the non-driver, genotypic fitnesses equal *w_CC_* =*w_AC_* =*w_AA_* =1, and *w_BC_* =*w_AB_* =1 − *s_h_*. Assuming HWE, a recessive fitness cost to drive, and assuming that the *A*/*B* locus is at its equilibrium frequency, the change in frequency of a rare drive modifier is

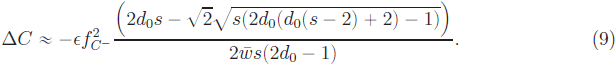

For all values of interest (0 < *s* < 1, 0.5 < *d*_0_ < 1), the change in frequency a rare *C* allele is positive when it decreases drive (i.e. *ϵ* < 0, corresponding to the A^−^ model, above), a result which holds qualitatively for a common *C* allele, as well (File S2).

### Model 6. Unlinked drive-modifier

For the unlinked model, we introduce another locus where drive is modified in *A*/*B* females fertilized by *M* allele, while the wild-type *L* allele does not influence drive. Assuming HWE and linkage equilibrium, the change in frequency of a rare unlinked, sperm-acting drive modifier is

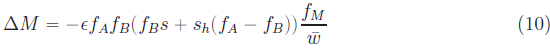

Thus, a rare drive suppressor (ϵ < 0) will spread so long as the fitness cost of the driver does not display over- or under-dominance.

**Model 6′. Unlinked Female Acting Drive-modifier**

The dynamics of a female-acting drive modifier are comparable to those describing a sperm-acting drive modifier. Assuming Hardy-Weinberg and linkage equilibrium, the change in frequency of a rare, unlinked, female-acting drive modifier is

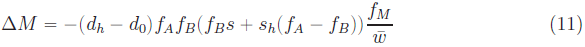

When when drive-modification is dominant (*d_h_* =*d*_1_ =*d*_0_ + ϵ), Equation 11 is equal to Equation 10. However, if female drive suppression is less than fully dominant, sperm-acting drive suppressors are more efficacious when rare than are female-acting suppressors, and are therefore more likely to spread.

## Supplementary Material

**File S1:** Transmission rules. We detail the frequency of offspring genotypes produced by each possible mating for Models 1-6 in the tabs of this Excel spreadsheet.

**Files S2A & S2B:** A Mathematica file (FileS2A) and a PDF of this file (File S2B) in which we derive analytical results for models 1-6 and 6−.

**File S3:** Exact approach. The R Script used for exact recursions for all models, including cases with inbreeding.

**File S4:** Variation in critical time-points during female meiosis across taxa (adapted from Masui (1985)).

**File S5:** An R object containing the phylogeny and raw data used to generate Figure S1.

**File S6:** The R Script used to generate Figure S1. This requires that File S5 is loaded into the R environment.

### Supplementary-Figures

**Figure S1:**
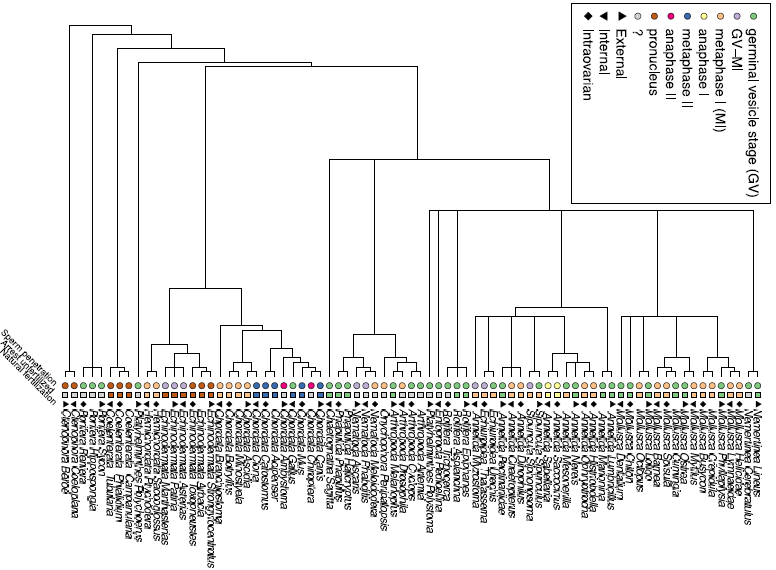
The phylogenetic distribution of the female meiotic arrest. The first two columns of symbols gives the stage of the ooycte when the sperm penetrates and the stage at which it arrests if unfertilized. The third column of symbols shows the site for natural fertilization. These columns of phenotypic data were extracted from Table 1 of Masui (1985). The tree was extracted from the Open Tree of Life project. The raw data table, the phylogeny/supporting R objects, and the script to do this is are included in the supplement (Files S4-S6).

**Figure S2:**
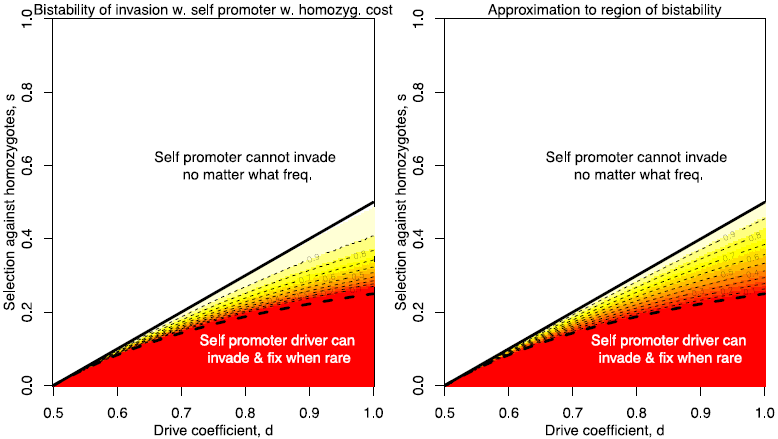
Invasion analysis for a self-promoting female meiotic drive allele with recessive costs (selection coefficient s), showing the region of bistability. The colors, and the thin dashed contours, indicate the frequency the allele must reach, *f*^*^ in order to invade the population (note that these alleles reach fixation conditional on invading). In the white area, the allele cannot invade, in the solid red area the allele can invade and fix when rare. In the left panel we show the results obtained by a grid search using the recursion, on the right we show the approximation obtained assuming that HWE holds.

**Figure S3:**
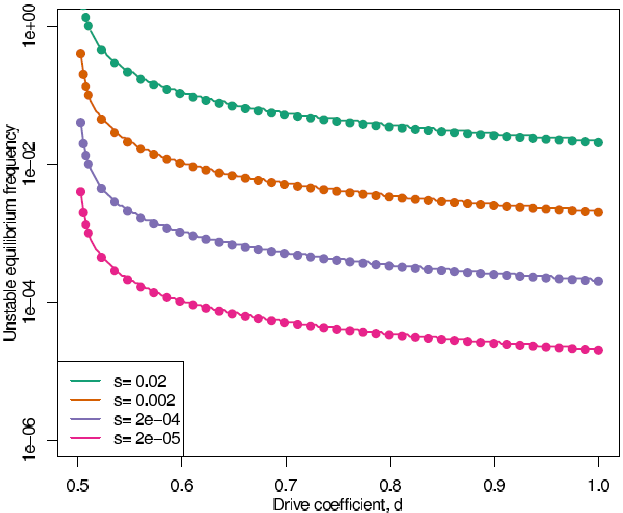
The unstable equilibrium frequency for a self-promoting female meiotic drive allele with an additive cost (*s_h_* =*s*/2) as a function of the drive parameter. The solid line shows results obtained using the recursion, the dots our approximation given by Equation (5).

**Figure S4:**
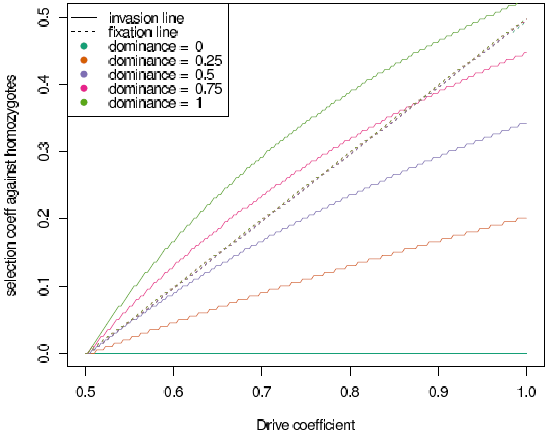
Exact results of invasion analysis of an allele whose effect on female meiosis is mediated by the genotype of the fertilizing male. A heterozygous female transmits the *B* allele with probabilities 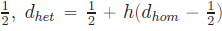, if she mated with a n *AA*,*AB*,or *BB* male, respectively. The allele suffers a recessive fitness cost *s*. The four panels correspond to different dominance relationships. In the parameter space below the invasion (solid) line the self-promoter driver can invade. In the parameter space below the fixation (long dash) line the self-promoter can fix. In the last two panels the invasion line is above the fixation line and so the allele can be maintained as a polymorphism in that thin slice of parameter space between the two lines. In the final panel we show the fixation line (small dashes) as predicted by our HWE approximation ((*d_he_t* − 1/2)/*d_het_*) see the appendix for more details.

Figure S5: Download from [https://brandvainlab.files.wordpress.com/2014/12/figs5.pdf]Evolution of a self-promoter and standard driver with variable levels of inbreeding (modifying the selfing rate from 0 to 0.9, in 0.1 increments). Assuming that the fitnesses of drive homozygotes and heterozygotes are 1 − *s* and 1, respectively. Boundary conditions for the invasion (solid lines) and fixation (dotted lines) of self-promoting (red) and standard (black) meiotic drivers, with drive coefficient, *d*. We derived these conditions from the simulation in File S3.

